# Paracrine behaviors arbitrate parasite-like interactions between tumor subclones

**DOI:** 10.1101/2020.12.14.422649

**Authors:** Robert J. Noble, Viola Walther, Christian Roumestand, Urszula Hibner, Michael E. Hochberg, Patrice Lassus

## Abstract

Explaining the emergence and maintenance of intratumor heterogeneity is an important question in cancer biology. Tumor cells can generate considerable subclonal diversity, which influences tumor growth rate, treatment resistance, and metastasis, yet we know remarkably little about how cells from different subclones interact. Here, we confronted two murine mammary cancer cell lines to determine both the nature and mechanisms of subclonal cellular interactions *in vitro.* Surprisingly, we found that, compared to monoculture, growth of the ‘winner’ was enhanced by the presence of the ‘loser’ cell line, whereas growth of the latter was reduced. Mathematical modeling and laboratory assays indicated that these interactions are mediated by the production of paracrine metabolites resulting in the winner subclone effectively ‘farming’ the loser. Our findings add a new level of complexity to the mechanisms underlying subclonal growth dynamics.

## Introduction

Considering tumors as complex ecosystems has led to the notion that diverse cancer cell subclones engage in heterotypic interactions reminiscent of those that operate in organismal communities (Heppner, 1984; Merlo et al., 2006; Axelrod et al., 2006; Tabassum and Polyak, 2015). Mutually negative interactions are thought to be ubiquitous in cancer (Nowell, 1976; Greaves & Maley, 2012). As in classic ecosystems, cancer cells compete for nutrients and space, and competition between emergent subclones gives rise to complex temporal and spatial dynamics of tumor composition and growth (Tabassum and Polyak, 2015; Freischel et al. 2021). Positive ecological interactions (mutualism and commensalism) have been observed in cancer models in mice (Calbo et al., 2011; Cleary et al., 2014) and in drosophila (Ohsawa et al., 2012). In these cases, one subclone acquires new abilities, such as the capacity to grow or metastasize, only in the presence of another subclone, resulting in the tumor as a whole progressing towards a more aggressive phenotype. In contrast, the prevalence within tumors of asymmetric interactions such as amensalism, parasitism and facilitation remains an open question. Defining the mechanisms of tumor ecology is essential for a better understanding of cancer progression and may lead to novel therapeutic strategies (Gatenby and Brown, 2017; Maley et al., 2017).

To gain insight into molecular and cellular events related to ecological interactions between cancer subclones, we took advantage of a model described over three decades ago, based on two closely related murine cancer cell lines derived from a single spontaneous mouse mammary tumor (Dexter et al. 1978; Miller et al., 1988). When cultured separately, the two cell lines have similar growth rates, yet in co-culture one cell line (the ‘winner’) expands at the expense of the other (the ‘loser’). Our careful re-examination of this model, combining experiments with mathematical modelling and parameter inference, indicated that the cellular behaviors of the two subclones are surprisingly sophisticated. Both cell lines produce paracrine metabolites that boost proliferation of the winner and also decrease the growth rate of the loser. Our results thus unveil a type of facultative parasitic behavior of the winner subclone. We further identified beta-hydroxybutyrate and lactate as metabolites that contribute to these phenotypes and characterized their modes of action. We discuss our results in the context of how previously underappreciated ecological interactions may contribute to the complexity of tumor growth dynamics.

## Results

### 4T07 cells have a “winner” phenotype

Two cell lines derived from a single mouse mammary carcinoma - 168 and 4T07 cells - have similar growth rates when cultured individually, yet the 4T07 clone displays a dominant phenotype when grown together, either in cell culture or in orthotopic allografts *in vivo* (Miller et al., 1988). Several hypotheses to account for this interesting behavior had been tested in the original work, but the precise mechanism behind these competitive interactions has so far not been identified.

We began by verifying that in our hands the lines maintain their competitive characteristics. To facilitate lineage tracing we first generated lines stably expressing GFP, the expression of which did not alter cell growth (Figure 1A). Next, we followed growth characteristics of 4T07 and 168FARN cells, the latter being a drug-resistant derivative of the original 168 clone (Aslakson et al., 1991), in a continuous culture for 3 weeks. The cells were seeded as 1:1 mix at a density that allowed them to reach confluence within 3-4 days, at which point they were harvested and re-seeded in a new well at the original density. Remaining cells were analyzed by flow cytometry to determine the proportion of GFP expressing clones in the expanding population.

**Figure 1.**
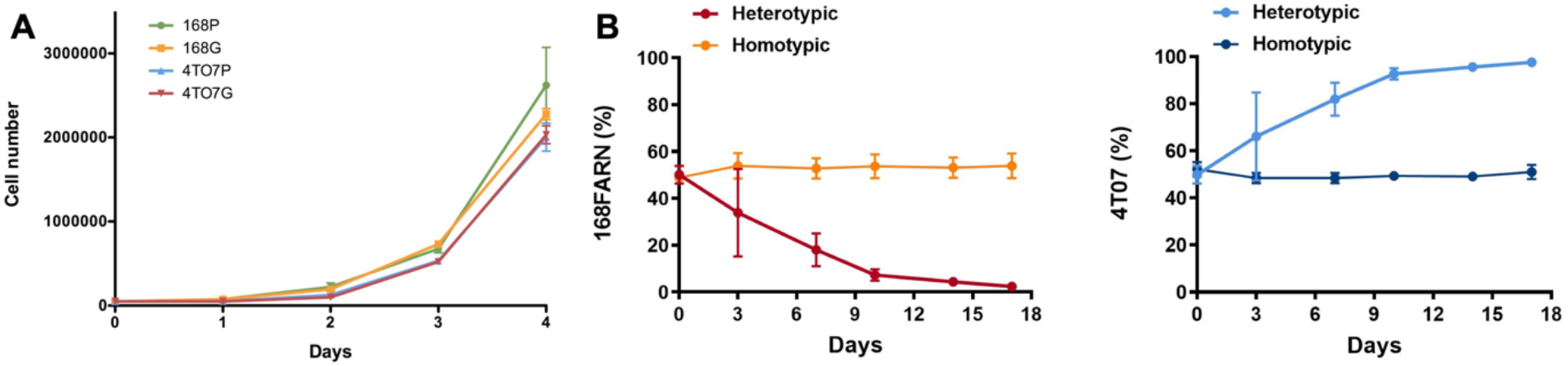
Mutual impacts on subclonal growth. **A:** 168FARN and 4T07 parental cells were transduced either with an empty retroviral vector (168P and 4T07P) or with labelled with a GFP-encoding retrovirus (168G and 4T07G). Cells were seeded in triplicate in 6-well plates at a density of 50 000 cells/well and cultured for the indicated times before harvesting and counting. **B:** 10^5^ cells were seeded at a 1:1ratio in homotypic (parental and GFP expressing derivative of the same cell line) or heterotypic (different cell lines, one expressing GFP) co-cultures and harvested and replated at the initial densities (10^5^ cells/plate) at indicated times. The ratios of GFP-labelled to unlabelled cells were estimated by flow cytometry. The results represent data from 3 independent experiments and are shown as mean +/− SEM.

The homotypic co-culture (same line with and without GFP) confirmed that GFP has no impact on cellular proliferation (Figure 1B and Figure 3B). In contrast, heterotypic co-culture conditions (two different lines, one expressing GFP) revealed the dominance of the 4T07 clone (Figure 1B and Figure 3B).

These results confirm the originally described ecological interaction between the clones: 4T07 gradually dominates the culture while the 168FARN cells become scarce within 15-17 days. Importantly, the dominant phenotype is independent of the starting ratio between the two cell lines (Supplementary Figure 1A and B).

### Co-culture alters the proliferation rates of both “winner” and “loser” cells

As originally discussed for the two clones under study (Miller et al., 1988), the expansion of a single clone in co-culture could be due to alterations in cell death or changes in the proliferation rates of either or both clones. We measured apoptosis in the loser 168FARN clone and found identical, very low levels of cell death under homotypic and heterotypic conditions (Supplementary Figure 2A). Next, we used time-lapse microscopy to assess the growth dynamics of both clones in continuous culture. The cells were seeded at a density that allowed reaching confluence in 4 days and were photographed every 45 minutes for the last 3 days. We measured the overall pixel intensity for each frame (Figure 2A) as a proxy for the growth rate of the fluorescently tagged cell line. This analysis revealed that under co-culture conditions, the growth rate of 168FARN decreased, whereas that of 4T07 increased relative to mono-cultures. To test whether increased net growth of the winner population is due to the alteration of proliferation, we estimated the proportion of cells in the S phase of the cell cycle by performing pulse-chase EdU staining. The results presented in Supplementary Figure 2B confirmed that heterotypic co-culture gave rise to significant decrease in cells actively replicating DNA for the loser clone and a significant increase in the winner clone. Overall, these results suggest that the dominant phenotype displayed by the winner cells in co-culture can be explained by changes in proliferation that operate in opposing directions on the winner and the loser cells.

**Figure 2.**
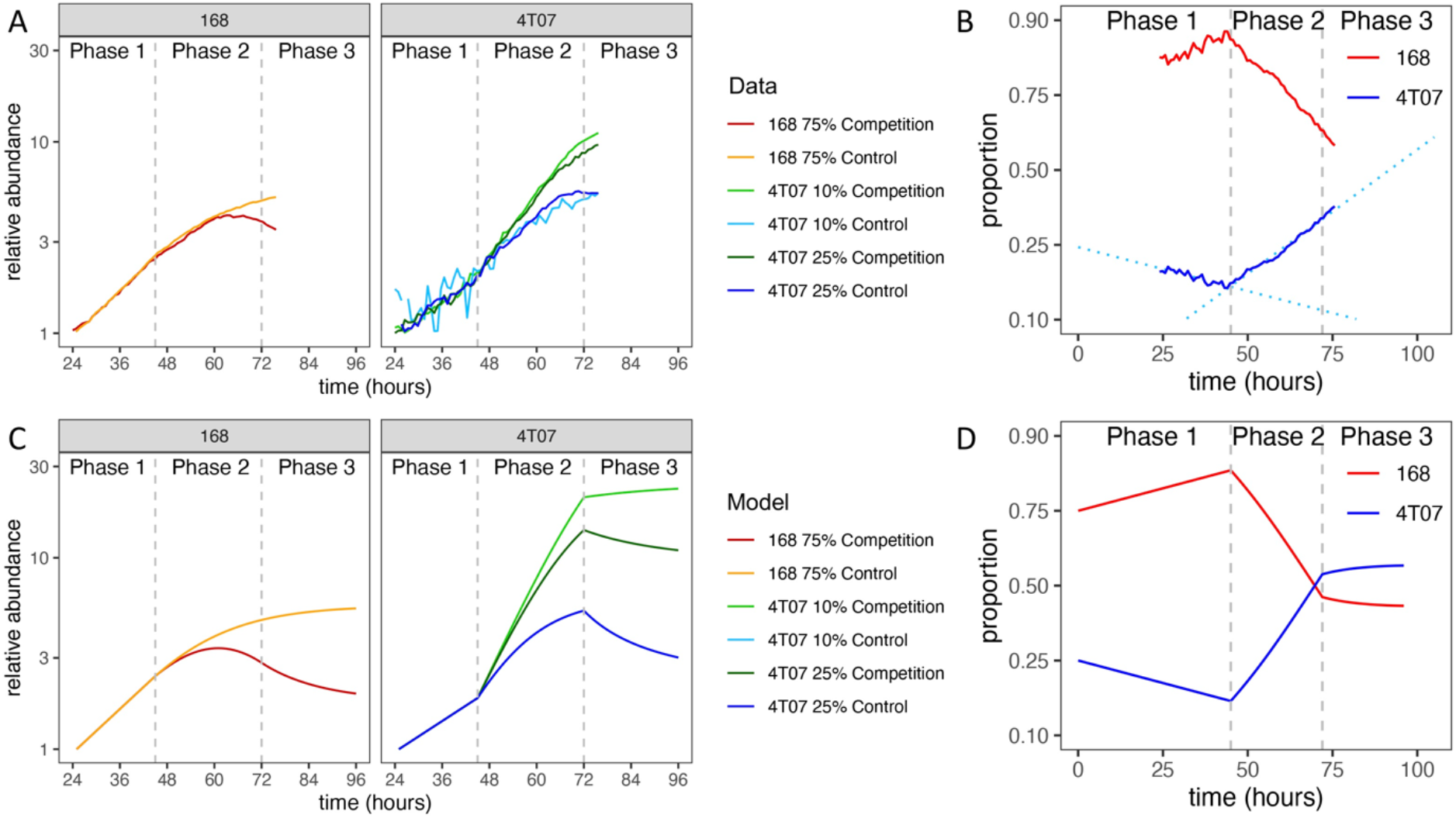
Normalized growth curves of homotypic and heterotypic mixes of subclones. **A:** The GFP fluorescence of the labeled subclone was measured by time-lapse microscopy. Cultures were seeded with 10^5^ cells per well. Log-transformed data were normalized by fitting regression lines and dividing by the inferred value at 24 hours. Vertical dashed lines mark the start of phase 2 (45 hours) and phase 3 (72 hours). **B: Frequency dynamics.** Curves obtained by combining the results of two competition experiments: one with labelled 4T07 and the other with labelled 168. The initial 4T07 proportion was 25% in both cases. The vertical axis is logit-transformed so that the slope of each curve is equal to the difference in net growth rates at the corresponding time (see Methods). Dotted regression lines are shown to draw attention to the change of slope. **C: Normalized growth curves according to mathematical model with parameter values inferred from data.** The model is described in Methods and parameter values are given in Table 1. **D: Frequency dynamics according to mathematical model with parameter values inferred from data.**

### Mathematical modelling and inference of evolutionary parameter values

To gain further insight into the ecological interactions between the winner and loser cell types we turned to mathematical modelling. Examination of the growth curves revealed two distinct phases of evolutionary dynamics (Figure 2A and 2B). In phase 1, from 0 to 45 hours, the two cell types grew exponentially in both homotypic and heterotypic cultures, and the growth rate of 168 was higher than that of 4T07. This first phase can be regarded as a transition period before the cells start altering and responding to their new environment. By contrast in phase 2, from 45 to 72 hours, the growth curves were strongly affected by interactions within and between the two cell types, and 4T07 grew faster than 168. We therefore assumed a mathematical model with exponential growth in phase 1 and a transition to density-dependent competitive Lotka-Volterra-type dynamics in phase 2.

By fitting our model to the homotypic growth curves, we inferred the values of the phase 1 and phase 2 growth rates and the within-type interaction parameters (Methods). To infer the between-type interactions, we used additional data from 72-hour competition assays, covering a wide range of initial ratios of the two cell types. Although this latter data set comprises only the initial and final proportions (at the beginning of phase 1 and the end of phase 2), we were able to infer the proportions at the beginning of phase 2 by adjusting for the exponential growth of both types during phase 1. We then used these inferred proportions and our previously inferred parameter values to estimate the remaining interaction parameters (Methods). The resulting model gives a good fit to the competition assay data (Figure 3A, first column) and is consistent with heterotypic time-lapse data not used for parameter inference (Figure 2; Supplementary figure 6).

**Figure 3.**
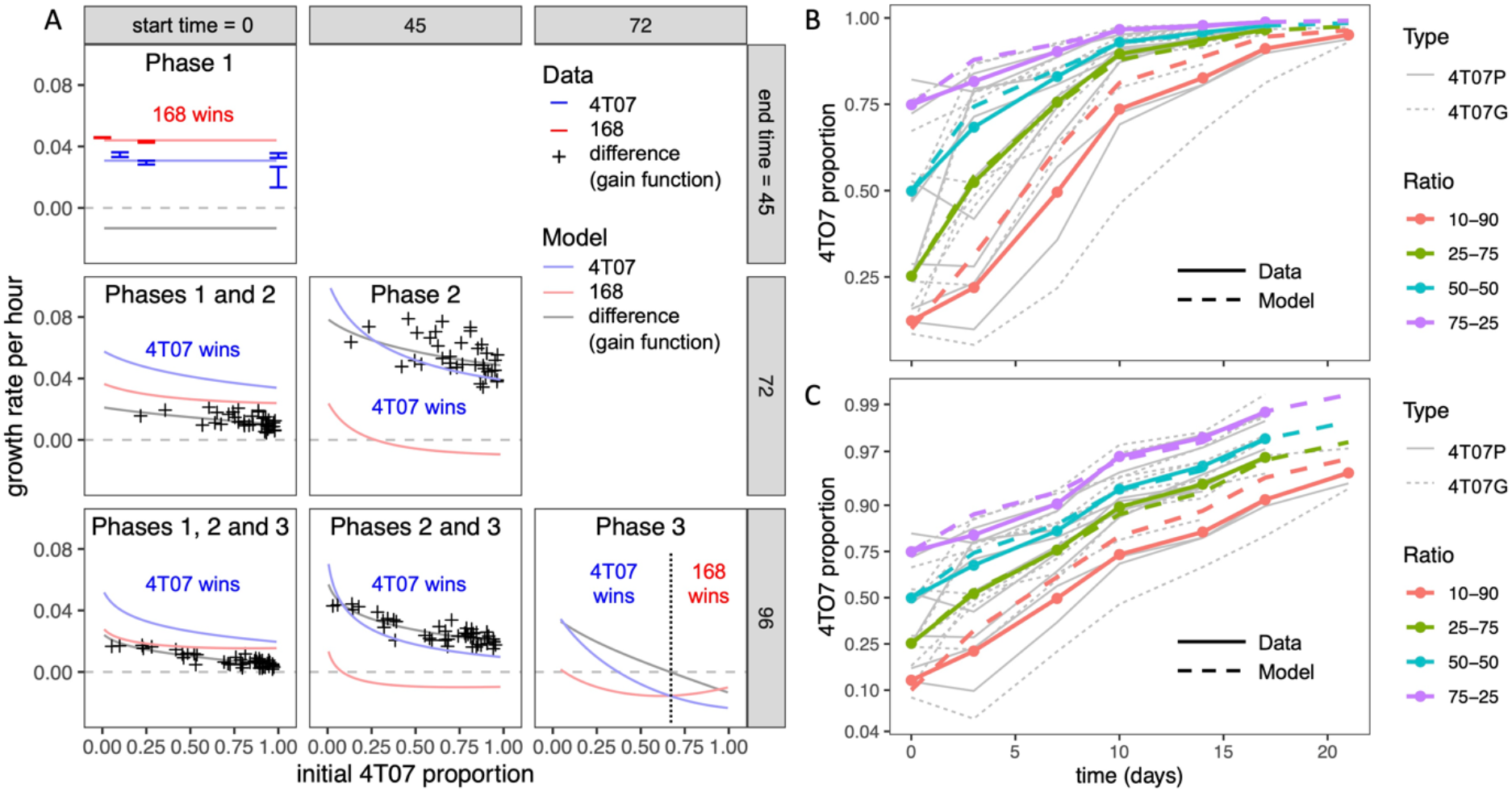
Mean net growth rate differences according to mathematical model and experimental data. **A: Inferred mean net growth rates and mean net growth rate differences (gain functions) over different time periods, corresponding to different phases within competition assays.** Columns correspond to different start times and rows to different end times of the phase(s) under consideration. For example, the centre panel labelled ‘Phase 2’ corresponds to the period between 45 and 72 hours. The initial 4T07 proportion (horizontal axis) is measured at the start of the respective period and the growth rate (vertical axis) is averaged over the period. Phase 1 data are from time-lapse microscopy. Other data points in the first column are from serial competition assays, such that each point corresponds to the slope of a thin grey line in **B**. Data points in the middle column are obtained from the competition assay data by adjusting for exponential growth during phase 1 (see Methods). Curves are the results of our mathematical model (Methods) with parameter values inferred from data (Table 1). **B: 4T07 frequency dynamics across serial competition assays.** Thick solid lines are averaged data (means of replicates with similar initial 4T07 proportions) and thick dashed lines are results of our mathematical model with parameter values inferred from data. Thin grey lines are data for individual experiments. A total of 10^5^ cells were seeded in co-cultures and harvested and replated as indicated. 4T07 parental cells were transduced either with an empty retroviral vector (4T07P) or labelled with a GFP-encoding retrovirus (4T07G). The ratios of GFP to unlabelled cells were estimated by flow cytometry. **C: Logit-transformed 4T07 frequency dynamics.** This panel shows the same data as **B** but with a logit-transformed vertical axis so that the slope of each curve is equal to the mean net growth rate difference (the gain function, as described in Methods and Supplementary figure 7).

The inferred parameter values (Table 1) imply that during phase 2, 4T07 has a large negative effect on both itself and on 168, consistent with 4T07 producing a harmful diffusible factor. The negative effect of 168 on itself is only about half as large, and 168 has approximately zero net effect on the growth of 4T07. This suggests that ubiquitous negative effects of 168 on 4T07 (e.g., likely due to waste products and competition for resources) are offset by positive effects, such as due to a beneficial diffusible factor. Also, during phase 2, the intrinsic growth rate of 168 (that is, the inferred growth rate before accounting for cell-cell interactions) is approximately 30% lower than that of 4T07, consistent with the conventional hypothesis that producing beneficial factors is costly. This disadvantage is offset by 168 having an approximately 30% higher carrying capacity (defined as the upper limit of the homotypic population size). Over phase 2, or any longer period that includes phase 2, the inferred net growth rate of 4T07 (that is, the growth rate after accounting for cell-cell interactions) is invariably higher than that of 168, which means 4T07 will come to dominate numerically, no matter their initial frequency.

**Table 1.** Mathematical model parameter values inferred from data. The interaction terms *a*, *b*, *c* and *d* are relative to population size, which is, in turn, relative to initial population size.

| Parameter | Phase(s) | Inferred value | Interpretation |
| --- | --- | --- | --- |
| $r_{L,1}$ | 1 | 0.044 | 168 growth rate in phase 1 (per hour) |
| $r_{W,1}$ | 1 | 0.031 | 4T07 growth rate in phase 1 (per hour) |
| $r_{L,2}$ | 2 and 3 | 0.073 | 168 intrinsic growth rate in phase 2 (per hour) |
| $r_{W,2}$ | 2 | 0.102 | 4T07 intrinsic growth rate in phase 2 (per hour) |
| $r_{W,3}$ | 3 | 0.04 | 4T07 intrinsic growth rate in phase 3 (per hour) |
| $a$ | 2 and 3 | -0.004 | Density-dependent effect of 168 on 168 |
| $b$ | 2 and 3 | -0.010 | Density-dependent effect of 4T07 on 168 |
| $c$ | 2 and 3 | 0.000 | Density-dependent effect of 168 on 4T07 |
| $d$ | 2 and 3 | -0.008 | Density-dependent effect of 4T07 on 4T07 |
| $K_L = -r_{L,2}/a$ | 2 and 3 | 17 | 168 carrying capacity, relative to initial population size |
| $K_W = -r_{W,2}/d$ | 2 | 13 | 4T07 carrying capacity, relative to initial population size |
| $\beta = b/a$ | 2 and 3 | 2.4 | Effect of 4T07 on 168, relative to effect of 168 on 168 |
| $\gamma = c/d$ | 2 and 3 | 0.0 | Effect of 168 on 4T07, relative to effect of 4T07 on 4T07 |

Since we also conducted 96-hour competition assays, we were able to infer the population dynamics during a third phase (72-96 hours). For every initial ratio of the two cell types, the growth rate difference (also known as the gain function) was on average lower in the 96-hour than in 72-hour competition assays (Supplementary figure 5). Moreover, this difference did not depend on the initial ratio, which implies it was not caused by a change in interaction parameters. A parsimonious way to account for this effect is to assume a reduction in 4T07’s intrinsic growth rate during phase 3, as would be expected to result from starvation and/or the build-up of toxic waste products. Making this adjustment to our model indeed produces a better fit to the competition assay data (Figure 3A, middle column; Figure 3B and 3C). The predicted dynamics are shown in Figure 2C and 2D.

Finally, having inferred all the evolutionary parameter values, we calculated net growth rates of the two cell types, averaged over different time periods. Over any period that includes phase 2, our model predicts that the net growth rate of both cell types will decrease non-linearly with increasing initial 4T07 frequency (pink and blue curves in Figure 3A). However, the net growth rate of 4T07 decreases faster than that of 168, which is why the gain function (grey curve in Figure 3A) also decreases. In phase 3, if the initial proportion of 4T07 is high (above 70%), then 168 has a higher net growth rate than 4T07, but in this case both of the inferred net growth rates are negative. Overall, the interactions are effectively equivalent to those of a parasite and its host, such that the ‘loser’ 168 suffers from the presence of the ‘winner’ 4T07, while also enhancing the winner’s fitness.

### β-hydroxybutyrate secreted by the loser clone stimulates winner clone proliferation

To identify the molecular mechanisms at the basis of the altered growth of winners and losers when in co-culture, we first focused on the increase in proliferation rate of 4T07 cells. Heterotypic culture experiments performed at low cell density suggested that the dominant effect did not require extensive cell-cell contacts (Supplementary Figure 3). We reasoned that a soluble factor secreted by 168FARN could induce a proliferation boost in 4T07. To test this hypothesis, we collected conditioned media from each line cultured for three days and used each medium separately to grow 4T07 for an additional 24 hrs. As controls, we either left the 4T07 medium after the three days of conditioning or replaced it with fresh medium. The results shown in Figure 3A confirm our hypothesis: the medium conditioned by 168FARN induced a significant increase in 4T07 proliferation. Importantly, this effect was not due to differences of medium exhaustion by the two cell lines, since the addition of fresh medium did not boost 4T07 proliferation.

Since our data strongly suggested that a soluble factor originating from 168FARN accounted for the increase in 4T07 proliferation, we next sought to define its molecular nature. First, we separated the 168FARN-conditioned medium into high and low MW fractions with a 3 KDa molecular cutoff column. The low MW fraction contains mainly metabolites while the high one is enriched in proteins. After complementing each fraction, respectively, with 10% serum or with DMEM to obtain full media conditioned with either low or high MW secretomes, we used them in a proliferation assay as in Figure 4A. The results (Figure 4B) of this series of experiments unambiguously identified the low MW fraction of the 168FARN-conditioned medium as the source of the pro-proliferative factor. To further explore its identity, we employed nuclear magnetic resonance spectroscopy to compare the composition of low MW fractions prepared from fresh medium and from the 168FARN- and 4T07-conditioned ones (Henke et al., 1996). Two major peaks specific for the conditioned media corresponded to a very strong signal for lactate secreted by 4T07 cells, and a significant increase in a peak identified as β-hydroxybutyrate in the 168FARN-conditioned medium (Figure 5A). β-hydroxybutyrate (BHB) is a ketone body mainly produced by the liver after long fasting periods and which is used by different tissues as a source of carbon to supplement the lack of glucose (Newman and Verdin, 2017). In addition, β-hydroxybutyrate is also produced by other cell types, such as adipocytes or cancer cells (Grabacka et al., 2016; Huang et al., 2017; Wang et al., 2017). To confirm the NMR-based identification of the β-hydroxybutyrate peak, we employed an enzymatic assay to measure β-hydroxybutyrate concentration in conditioned media from 4T07 and 168FARN (Figure 5B). The results were in perfect agreement with the NMR analysis: β-hydroxybutyrate production is significantly higher in the loser than in the winner cell clone. To test whether this metabolite was indeed responsible for the increased proliferation of 4T07, we next complemented the medium of exponentially growing 4T07 cells with purified β-hydroxybutyrate. As shown in Figure 5C, β-hydroxybutyrate increased the 4T07 proliferation rate to a level comparable to that obtained with the 168-conditioned medium. We thus conclude that loser cells increase the winner’s growth rate through the secretion of β-hydroxybutyrate.

**Figure 4.**
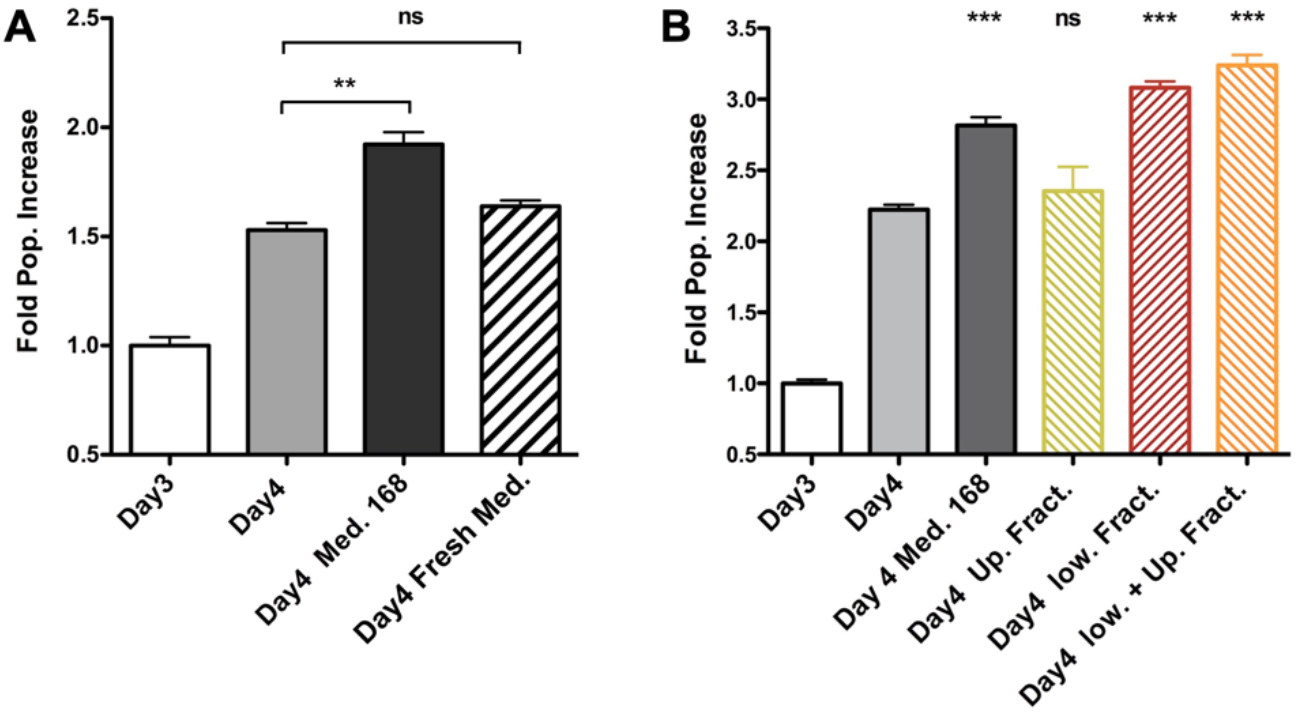
Soluble factor secreted by 168FARN cells accelerates proliferation of the 4T07 cells. **A:** 4T07 cells were grown for 3 days at which point their medium was either left unchanged, or replaced by either 168FARN-conditioned medium or fresh medium, as indicated. Cells were collected 24 hrs later and counted. Cell numbers at day 3 were arbitrarily set at 1 in order to include the data from 3 independent experiments. **B:** The experiment was performed as in **A.** but the medium conditioned by 168FARN cells was fractionated by membrane ultrafiltration with a 3 KDa molecular cutoff. After complementing the low and the high MW fractions, respectively, with 10% serum and DMEM, the media were used to grow the 4T07 cells, as in **A**. The two fractions were also combined as a control. ns: not significant, * p<0.05, **p<0.01, ***p<0.001, all compared to Day 4 point.

**Figure 5.**
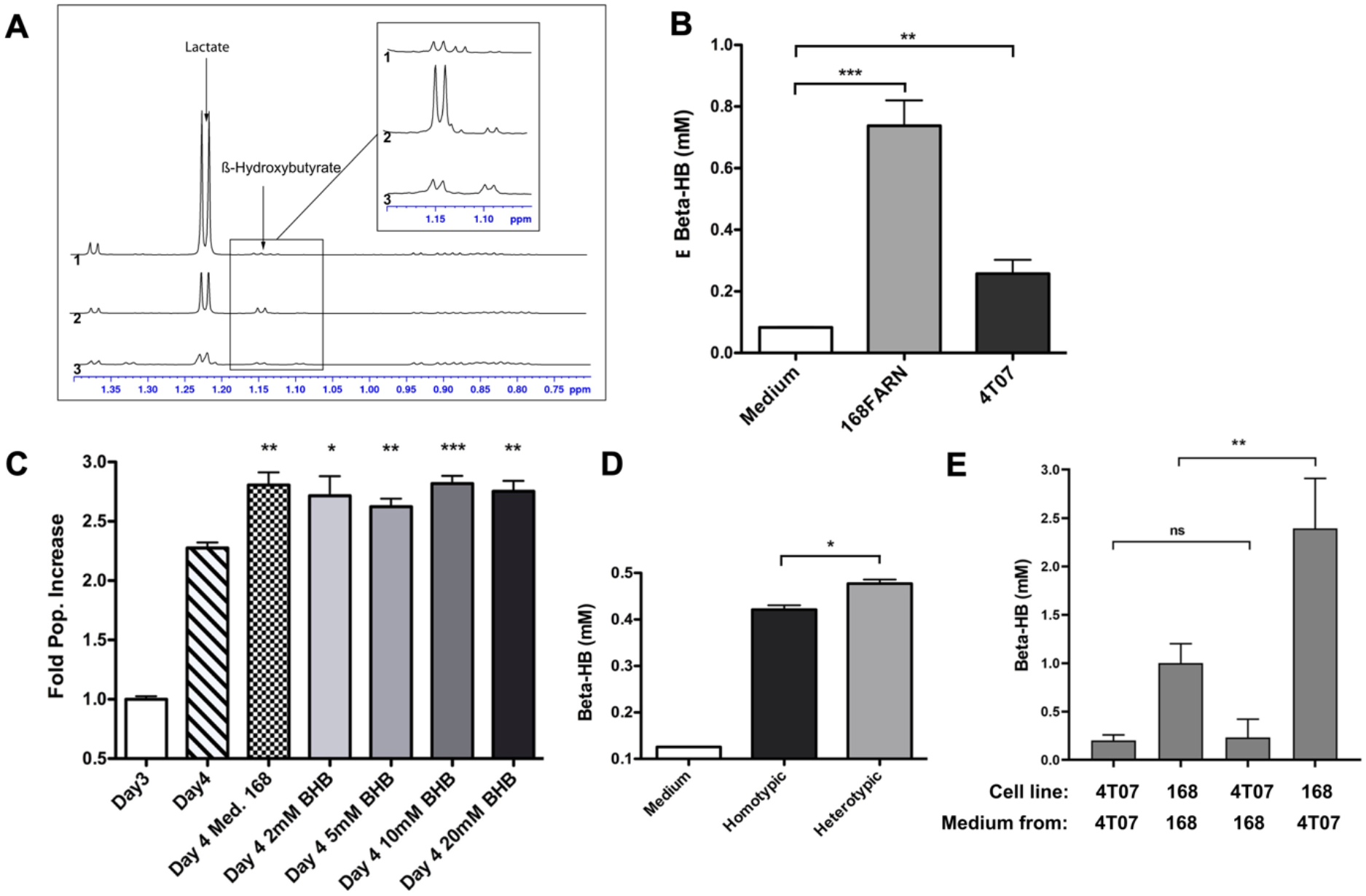
Identification of soluble metabolites altering the heterotypic growth dynamics. **A:** Superimposition of the high-field region of representative 1D proton NMR spectra recorded at 700 MHz, 293 K and pH7 on samples of culture media collected after growing 40T7 cells (1) or 168FARN cells (2) for 3 days or of fresh cell culture medium (3). The arrows indicate the characteristic resonance of Lactate and β-hydroxybutyrate. The insert displays a zoom in this spectral region, revealing the H-alpha resonance of the β-hydroxybutyrate. For all spectra, peak intensities have been normalized on the intensity of the DSS resonance added as internal reference. **B:** Concentration of β-hydroxybutyrate from fresh medium and from conditioned medium from 168FARN or 4T07 was quantified. **C:** Commercially available β-hydroxybutyrate at indicated concentrations was added to 4T07 cell culture at day 3 and the growth allowed to proceed for an additional 24 hrs. All points are compared to Day 4 point. **D:** 168FARN alone (homotypic) or in 1:1 co-culture with 4T01 cells were grown for 4 days and extracellular β-hydroxybutyrate was measured enzymatically as in 4B. **E:** 168FARN and 4T07 cells were cultured individually for 3 days. The medium was then replaced by the homotypic or heterotypic conditioned one, as indicated, and the culture allowed to continue for an additional 24 hrs. The β- hydroxybutyrate concentration was quantified at day 4. ns: not significant, * p<0.05, **p<0.01, ***p<0.001

### Presence of the winner clone stimulates β-hydroxybutyrate production by loser cells

After assessing β-hydroxybutyrate production in homotypic cell culture, we evaluated its secretion under heterotypic conditions. We grew 168FARN alone or together with 4T07 at a 1:1 ratio, maintaining the overall cell density constant. Surprisingly, despite the fact that under heterotypic conditions there are at least 50% fewer loser cells (which are the main producers of β-hydroxybutyrate, *cf.* Fig. 5B), the overall level of secreted β-hydroxybutyrate was higher than in the homotypic culture (Figure 5D). This suggests that either the presence of 4T07 increased the production of the metabolite by 168FARN or, alternatively, that it was 4T07 that produced more metabolite when grown in the presence of 168FARN. To distinguish between these hypotheses, we cultured both lines individually for three days, measured BHB concentration, and then exchanged the culture medium and quantified metabolite synthesis 24 hours later. The quantification of β-hydroxybutyrate produced over the last day (Day 4 BHB concentration minus Day 3 BHB concentration) shows that the 168FARN-conditioned medium had no effect on BHB secretion by 4T07 cells. In striking contrast, the production of the metabolite by 168FARN more than doubled under the influence of the 4T07-conditioned medium (Figure 5E). Thus, the winner cells stimulate the losers to produce a metabolite that boosts the former’s proliferation.

### Mechanism of β-hydroxybutyrate action

We next asked about the mode of action of BHB on the 4T07 cells. β-hydroxybutyrate can be imported by four monocarboxylate transporters of the SLC16A gene family, the expression of which varies in different cell types. We assessed the expression of each transporter by RT-QPCR and found that MCT2, MCT3 and MCT4 were barely expressed while MCT1 was highly expressed (Figure 6A) in 4T07 cells. This result suggests that MCT1 is likely responsible for the import of BHB in this cell line. Interestingly, we found that MCT1 is three times more expressed in 4T07 than in 168 cells (which, like 4T07, do not express the other MCTs - Supplementary figure 4A), suggesting that the winner cells are more efficient at taking up this metabolite than the losers (Supplementary Figure 4B). Finally, incubation of 4T07 with BHB upregulates MCT1, consistent with a positive feedback loop that could increase the transport of this ketone body into the dominant cell line (Supplementary figure 4C).

**Figure 6.**
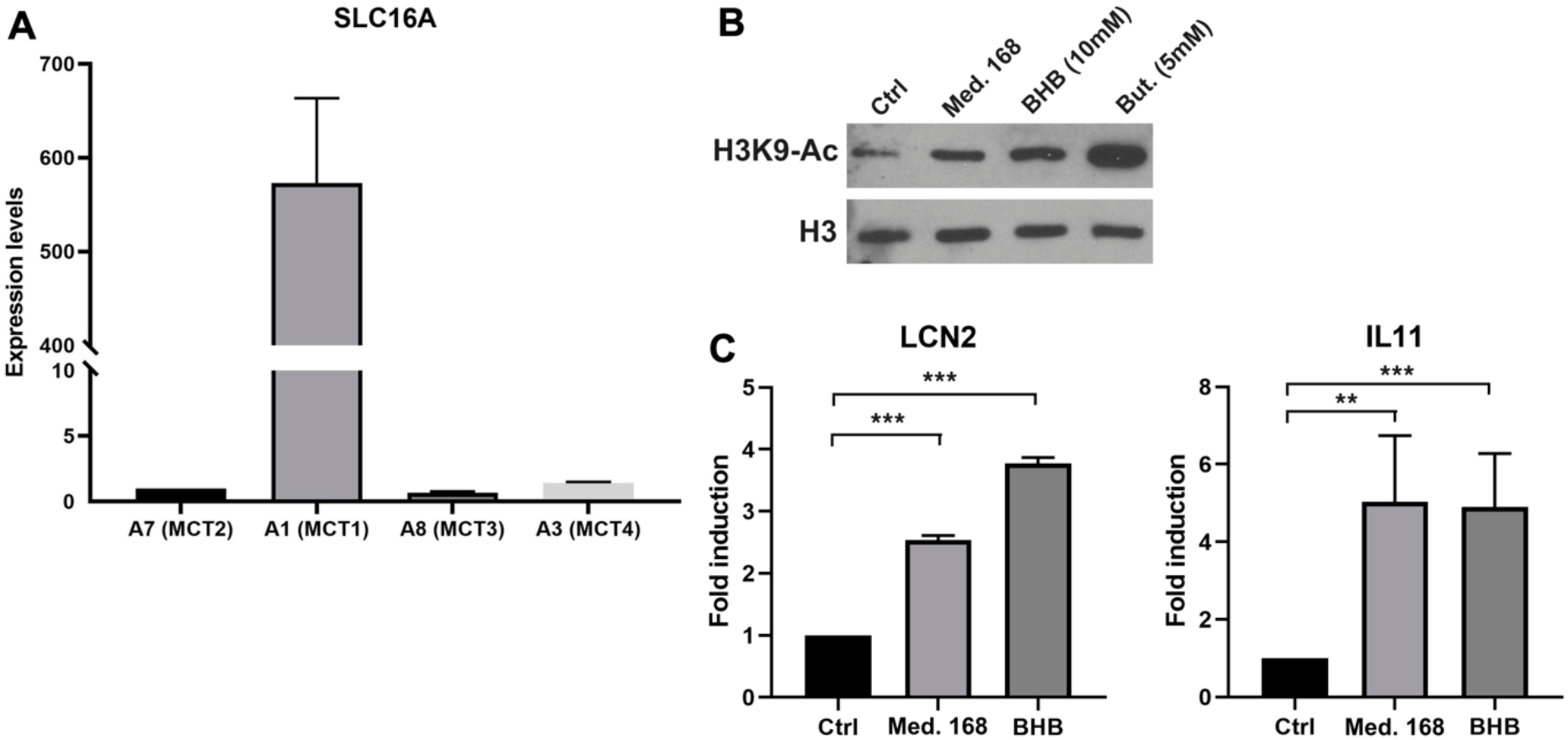
Extracellular β-hydroxybutyrate leads to increased H3K9 histone acetylation and altered gene expression in 4T07 cells. **A:** Expression levels of the slc16A family transporter genes in 4T07 were analyzed by RT-QPCR. Expression of HPRT served as normalization of the data. **B:** H3K9 histone acetylation was analyzed by immunblotting of extracts of 4T07 cells grown for 24 hrs in control, 168-conditioned medium or medium complemented with β-hydroxybutyrate or with butyrate, as indicated. Total histone 3 (H3) abundance served as normalization control.**C:** 4T07 cells cultured for 3 days were incubated for 8 hours with 4T07-(Ctrl) or 168-conditioned medium or purified β-hydroxybutyrate (10mM) added to fresh medium. Total RNAs were purified and subjected to RT-QPCR with specific primers for LCN2 and IL-11. **p<0.01, ***p<0.001.

β-hydroxybutyrate can be metabolized and used as a nutrient to replace glucose (Newman and Verdin, 2017). Experiments presented in Figure 3A show that fresh medium added at day 3 did not boost cell proliferation, suggesting that in this experimental setup the decrease in the carbon source is not a limiting factor for growth. It is thus unlikely that β-hydroxybutyrate is used as an energy resource to increase proliferation rate. β-hydroxybutyrate has previously been identified as an inhibitor of class I histone deacetylases (HDAC) that modulates the expression of genes involved in reactive oxygen species detoxification (Shimazu et al., 2013). Subsequently, another group found that adipocytes use β-hydroxybutyrate to modulate the expression of a subset of genes involved in the growth of breast cancer cells (Huang et al., 2017). We thus hypothesized that β-hydroxybutyrate might increase the growth rate of winners through the inhibition of HDACs, thereby modulating the expression of genes involved either in ROS detoxification or in the induction of pro-proliferative factors. In support of this idea, incubation of 4T07 cells either with 168FARN-conditioned medium or with purified BHB increased H3K9 acetylation, albeit to a lesser extent than butyrate, a bona fide HDAC inhibitor (Figure 6B).

While we could not detect in 4T07 cells any modification of expression of ROS detoxification genes reported to be regulated by β-hydroxybutyrate in other cellular models (Shimazu et al., 2013), both β-hydroxybutyrate and 168-conditioned medium led to significant transcriptional activation of interleukine 11 (IL-11) and lipocalin 2 (LCN2) (Figure 6C). Both genes have been previously described to promote cancer cell growth and to be regulated by β-hydroxybutyrate through its action on HDAC activity (Grivennikov, 2013; Huang et al., 2017; Yang and Moses, 2009). Thus, our data point to the molecular mechanisms involving direct proliferation signaling.

### Lactate secretion slows down loser cell proliferation

In addition to the positive effect of the 168FARN cells on the proliferation rate of the 4T07 clone, the data shown in Figure 2 indicate that the latter negatively influences the 168FARN growth dynamics. The NMR analysis highlighted strong lactate production (see Figure 5A). This is consistent with our observation of the media color change during culture of the two lines, indicating that the winner clone has a glycolytic type of glucose metabolism leading to a rapid medium acidification in culture. Because extracellular acidification can be detrimental for cell growth, we next asked if 168FARN were particularly sensitive to such growth conditions. We quantified medium acidification by seeding cells at different densities and measuring the extracellular pH after 3 days of culture (Figure 7A). As expected, we found that 4T07 cells acidify the medium faster and attain a lower pH during culture compared to 168FARN cells. Indeed, pH ranged from 6.94+/−0.005 (lowest density) to 6.79+/−0.003 (highest density) for the winner line and from 7.38+/−0.008 to 6.92+/−0.006 for 168FARN. To test whether 4T07-mediated extracellular acidification influenced 168FARN growth, we set up a proliferation assay for 168FARN cells grown in medium conditioned by the low and the high density grown 4T07 cells. To control for the effect of pH in the conditioned media, we included a treatment in which the medium from 4T07 was buffered at pH 7.0 by sodium bicarbonate. These experiments revealed that the medium from the low density 4T07 cells (pH 6.94) had no effect on 168FARN proliferation. In contrast, the medium from the high density 4T07 (pH 6.79) drastically decreased the 168FARN growth rate. Moreover, buffering the same medium at pH 7.0 restored the proliferative capacity of 168FARN culture (Figure 7B). We conclude that the loser clone is indeed highly sensitive to medium acidification. Taken together our data suggest that the decrease in the growth rate of 168FARN observed in heterotypic conditions is triggered by 4T07 mediated extracellular acidification.

**Figure 7.**
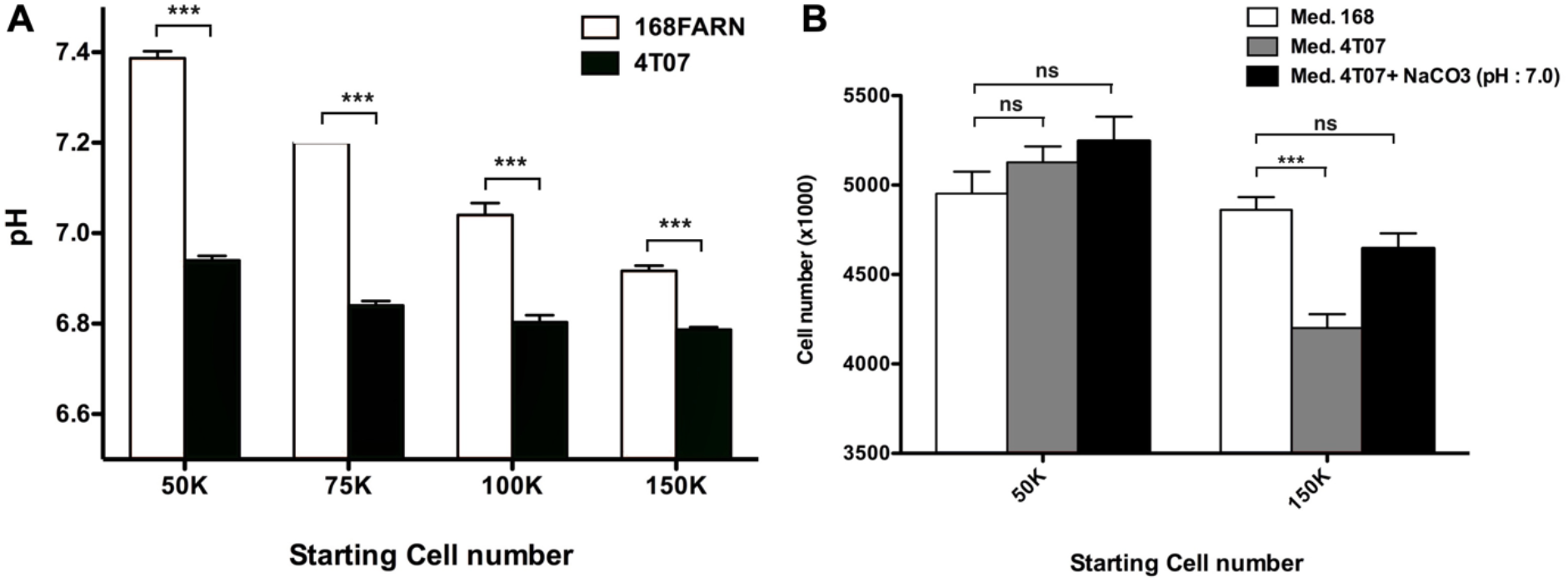
Impact of extracellular pH on the loser clone growth. **A:** 168FARN and 4T07 cells were seeded at the indicated initial densities in 6-well plates and cultured for 3 days. Culture media were removed, immediately covered with a layer of mineral oil to prevent oxidation and the pH was measured. **B:** 10^5^ 168FARN cells were grown for 3 days. Medium was then replaced by conditioned media from cultures grown at low or high density, as indicated. Where indicated, 5mM NaCO_3_ was used to buffer the 4T07 conditioned medium to pH7. 24 hours later cells were harvested and counted. Data are from three independent experiments conducted in triplicates. ns: not significant, ***p<0.001.

## Discussion

Heterogeneity is a ubiquitous feature of tumors that influences growth and metastasis, and thus the potential for therapeutic success. Ecological interactions between subclones are key to the emergence of this heterogeneity, yet only few empirical studies have characterized the nature of these interactions or their underlying mechanisms. These include commensal (Kaznatcheev et al. 2019; Farrokhian et al. 2020) and cooperative (Cleary et al. 2014) interactions *in vitro*, and how such interactions can drive tumor invasion (Chapman et al. 2014) and metastasis *in vivo* (Janiszewska et al. 2019; Naffar-Abu Amara et al. 2020).

Our study extends previous work (Robinson and Jordan 1989; Marusyk et al. 2014; Archetti et al. 2015) by demonstrating that two cell lines derived from the same tumor exhibit a sophisticated relationship, whereby one (the ‘winner’) effectively farms the population of the other (the ‘loser’). We further identified key metabolites (β-hydroxybutyrate and lactate) that regulate these interactions between the winning and losing clones. Similar to Archetti et al. (2015), we found that exploitative clonal interactions evolve through time, but whereas these authors observed a frequency-dependent change that could explain clonal coexistence, we were unable to detect this effect. Simple mathematical analysis within the framework of evolutionary game theory suggests that spatial heterogeneity might permit long-term clonal coexistence in our system (Methods).

Our *in vitro* experiments oversimplify the more complex interrelationships that predominate in spatially complex microenvironments, *in vivo*. That paracrine signaling is responsible for the effects we observed between winner and loser cell lines suggests that the spatial arrangement of these cells could be crucial to their growth and relative frequencies *in situ* (Archetti et al 2015). The effect of spatial structure would depend on the typical distance that secreted molecules travel through the complex tumor microenvironment. Our results indicate that areas of contact or close proximity between the two subclones will grow faster and therefore come to dominate spatially isolated populations, producing what is effectively a mixed 4T07-168FARN ‘phenotype’. The actual spatial arrangement of these two subclones in the original tumor is unknown, but the authors of the study originally isolating these cell lines note that they may represent only a small sample of the tumor’s diversity (Dexter et al., 1978). A growing body of evidence suggests that single, site-specific biopsies may be of little use in quantifying spatial heterogeneity, due to the multiscale (local, regional, metastatic) nature of tumor evolution (Amirouchene-Angelozzi et al., 2017). Computational modeling indicates that the range of cell-cell interaction and the mode of cell dispersal are crucial factors determining the pattern of intratumor heterogeneity and associated characteristics of tumor growth and evolutionary potential (Noble et al., 2020; Waclaw et al., 2015).

We find that the complex interactions between the 4T07 and 168FARN cells are governed by paracrine signaling emanating from both clones. This mechanistic conclusion differs from the original observations reported by Miller et al (Miller et al., 1988). Indeed, in the original publication the results concerning the inhibitory effect of 4T07 conditioned media on 168 cells were inconclusive. This apparent discrepancy could be due to slightly different culture conditions used in the two sets of experiments. Indeed, the medium acidification due to the lactate release by the 4T07 that is responsible for slowing down the growth of 168 cells reaches the required threshold value only after prolonged culture (3-4 days under our experimental conditions). It is thus possible that in the original report the culture time and/or the cell density were insufficient for the clear visualization of the paracrine effect of the winners on the losers. Moreover, Miller et al. did not investigate the paracrine effect exerted by the 168 on the 4T07 cells. Our results are the first to show the reciprocal effects of both cell lines on each other, thus highlighting the complexity of their mutual interactions.

We have identified a ketone body, β-hydroxybutyrate, which is produced by loser cells and acts to increase the growth rate of winner cells. Mechanistically, the competitive advantage afforded by β-hydroxybutyrate to the winner clone appears to be mediated through the HDAC-controlled activation of a genetic program that boosts its proliferation. Ketone bodies are small lipid-derived molecules, physiologically produced by the liver and distributed via the circulation to metabolically active tissues, such as muscle or brain (Newman and Verdin, 2017), where they serve as a glucose-sparing energy source in times of fasting or prolonged exercise. Recently, several studies reported that cell types such as adipocytes, intestinal stem cells or cancer cells originating from colorectal carcinoma or melanoma can also produce β-hydroxybutyrate (Cheng et al., 2019; Grabacka et al., 2016; Huang et al., 2017; Shakery et al., 2018). Our results identifying β-hydroxybutyrate as a signaling molecule involved in intra-tumoral clonal interactions fall into the general category of these novel roles for ketone bodies in cell communication.

However, the link between ketone bodies and tumor development remains controversial. On the one hand, it was shown that ketonic diet slows down tumor development in brain cancer mice models (Poff et al., 2013, 2014). On the other hand, our results together with other recent data (Huang et al., 2017) suggest that β-hydroxybutyrate may favor breast cancer progression. One unexplored possibility to explain these contradictory observations is that this ketone body can be used differently by different cancer cell types, for example as a carbohydrate supply or as a HDAC inhibitor, ultimately leading to cancer-type and context specific response.

In our experimental model, β-hydroxybutyrate increases winner cells proliferation by activating a genetic program through HDAC inhibition. Among the genes we discovered to be activated by the ketone body, IL-11 is an interleukin that displays a pro-proliferative activity (Grivennikov, 2013). Interestingly, in a distinct breast cancer cell cooperation model, sub-clonal expression of IL-11 favours the expansion not only of cells that express it, but also of other cellular sub-clones (Marusyk et al., 2014). This suggests that IL-11 acting in either paracrine or autocrine fashion could lead, respectively, to cooperation or to competition between subclones, thus participating actively in the selection and evolution of tumor heterogeneity.

Overall, our experimental data therefore suggest a model in which the winner line stimulates the production of and benefits from a compound delivered by the loser line and, conversely, the loser is negatively influenced by the presence of winners through secretion of another compound.

We note that while in artificially maintained conditions of non-constrained growth (in culture) the losers are eventually eliminated, many additional selective pressures that may affect clonal fitness operate *in vivo*. These involve cellular response to physical cues due to crowding (Vishwakarma and Piddini, 2020) and interactions with the extracellular matrix (Lu et al., 2012) as well as response to signaling from the stroma, including its inflammatory and immune components (Quail and Joyce, 2013). These elements are expected to influence the outcome of the direct interactions between the tumoral clones and may change the nature of their ecological interaction from net exploitation (*in vitro*) to mutual benefit (*in vivo*). Future study should evaluate whether parasitic effects are observed *in vivo*, and determine the extent to which these cell-cell interactions mediate important tumor characteristics, including growth, drug resistance, and metastatic behavior.

## Acknowledgements

The authors wish to thank HTE ‘HetCoLi’ (HTE20161) and ITMO ‘Physique Cancer’ (CanEvolve PC201306) for funding. RN acknowledges support from the National Cancer Institute of the National Institutes of Health under Award Number U54CA217376. The content is solely the responsibility of the authors and does not necessarily represent the official views of the National Institutes of Health. CR was supported by the French Infrastructure for Integrated Structural Biology (FRISBI) (grant No. ANR-10-INSB-05). MEH thanks the McDonnell Foundation for funding (Studying Complex Systems research award 220020294). We are grateful to Emie Quissac and Yasser Kerboua for their excellent technical assistance, and to Artem Kaznatcheev for helpful conversations.

## Methods

### Cell culture

4T07 and 168FARN were a kind gift of Dr Robert Hipskind. All cell lines were cultured in Dulbecco’s modified Eagle medium containing 10% fetal bovine serum, 100 ng/mL streptomycin, and 100 U/mL penicillin at 37 °C with 5% CO2.

For co-culture experiments a mixture of GFP-labelled and parental cells (empty-vector transduced) cells were seeded at the final density of 10^5^ cells/well in 6-well plates, except where mentioned otherwise. Upon reaching confluence (3-4 days) they were harvested, diluted to the original density and replated. The remaining fraction was analyzed by flow cytometry.

### Immunoblot Analysis

Cells were lysed in boiling Laemmli buffer supplemented with protease inhibitors, then sonicated and complemented with DTT. Protein concentration was determined by BCA (Thermo Scientific) assay. Fifteen to twenty micrograms of total protein were loaded onto SDS-PAGE gels and transferred onto nitrocellulose membranes. The membrane was blocked with TBST (1× TBS with 0.1% Tween 20) + 5% milk at room temperature for 1 h and incubated with primary antibody and then with horseradish peroxidase (HRP)-coupled secondary antibody (Amersham, Piscataway, NJ). Activity was visualized by electrochemiluminescence. Antibodies used in this study are anti Histone H3 (Cell signaling Technology #9717) and anti- Acetyl-Histone H3 (Lys9) (Cell signaling Technology #9649).

### Reverse Transcription and Real-Time PCR

Total mRNA was isolated using a RNeasy mini kit (Qiagen, Germantown, MD, USA). Reverse transcription was performed with random hexamers and M-MLV Reverse Transcriptase (Invitrogen). Real-time PCR was performed in triplicates with LC FastStart DNA Master SYBR Green I on a LightCycler rapid thermal cycler system (Roche Diagnostics, Mannheim, Germany), according to the manufacturer’s instructions. Housekeeping gene HPRT was used for normalization. Primers sequences are available upon request.

### Time-lapse microscopy

Time-lapse microscopy was performed at 37 °C with 5% CO2, with images taken at 45-minute intervals using an inverted Zeiss Axio-Observer microscope. The images were processed and analyzed using ImageJ software.

### EdU staining

Cells were incubated with 10μM EdU for 2 hours, harvested and processed using the Click-iT™ EdU Alexa Fluor™ 647 Flow Cytometry Assay Kit (ThermoFisher Scientific #C10424) following manufacturer instructions. Labeled cells were then analyzed on a FACSCalibur flow cytometer using CellQuestPro software (BD Biosciences).

### Apoptosis quantification

To determine the percentage of apoptotic cells with externalized phosphatidylserine (PS), adherent and floating cells were collected and labeled with the Annexin V-Cy3 Apoptosis Detection Kit (Abcam, Cambridge, UK, #ab14143) according to the manufacturer’s instructions. Labeled cells were then analyzed on a FACSCalibur flow cytometer using CellQuestPro software (BD Biosciences).

### β-hydroxybutyrate quantification

β-hydroxybutyrate concentration was measured by an enzymatic kit (Sigma-Aldrich MAK041) following the manufacturer instructions. Briefly, β-hydroxybutyrate present in the culture medium was determined by a coupled enzyme reaction, resulting in a colorimetric (450 nm) product, proportional to the β-hydroxybutyrate concentration. The absorbance was measured on a spectrophotometer.

### Medium fractionation

In order to separate low molecular weight molecules from the conditioned culture medium, 5 to 10 ml were loaded on a Vivaspin Turbo 15 PES, 3,000 MWCO column (Sartorius VS15T91) and centrifuged at 4000G for 30 minutes following the manufacturer instructions. Both fractions were then used for subsequent experiments and RMN analysis.

### RMN analysis

NMR experiments were recorded at 293K and pH 7 on an AVANCE III BRUKER spectometer operating at 700 MHz (proton frequency), using a Z-gradient shielded TCI 1H-13C-15N cryoprobe. Fully relaxed 1D 1H spectra were aquired with the regular 1D NOESY, using 5s as relaxation delay. The samples consisted on 1.5 mL of cell media (fresh or conditioned by cell culture), lyophilized and dissolved in 500 μL of deuterated phosphate buffer (50 mM, pH 7). DSS (EURISOTOP©, final concentration: 0.5 mM) was added as internal reference for chemical shift referencing and as a concentration standard for spectra normalization. The assignment of the 1H resonances of the compound of interest in this study (Lactate, β-hydroxybutyrate) was based on chemical shifts reported on the litterature (1) and further confirmed using 2D [1H,1H] (TOCSY) and [1H-13C] (HSQC, HSQC-TOCSY) NMR spectroscopy.

### Statistical analysis

Experiments were repeated at least three times. Data are presented as mean ± SEM. An Independent Student’s t test was performed to analyze the assay results; a two-tailed Student’s t test was used to compare the intergroup differences. Significance was accepted for values where P≤0.05 (*), P≤0.01 (**), P≤0.001 (***).

### Overview of mathematical methods

Our aim is to determine the general nature of the evolutionary dynamics in a form that can be readily compared to other systems. Our mathematical approach is in the same vein as that of Kaznatcheev (2017) and Kaznatcheev et al. (2019) but with three important differences. First, our method can accommodate a smaller data set and is thus more economical because we mostly rely on measurements of initial and final proportions in competition assays, such as can be determined via flow cytometry, rather than extensive time-lapse image analysis. Second, whereas Kaznatcheev (2017) and Kaznatcheev et al. (2019) confine their analysis to exponential or logistic growth phases, we also examine phases in which cell populations exhibit non-logistic dynamics. Third, because we consider non-logistic growth phases, we use a density-dependent rather than a frequency-dependent model. Our method is generic and, in principle, can be adapted to any experimental evolution set-up with two competing populations of cancer cells, bacteria, or other entities.

### Definitions and mathematical relationships

We define the intrinsic growth rate as the exponential growth rate in the absence of interactions. In the Lotka-Volterra differential equations, this parameter is multiplied by the population size of the respective type. The intrinsic growth rate is the limit of the net growth rate as the population sizes approach zero (when interaction terms are negligible).

We define the net growth rate as the actual rate of change of the population size (i.e. the time derivative), which is the sum of the basic growth rate and interaction terms.

Supplementary figures 6 and 7 illustrate some of the mathematical relationships relevant to our methods.

### Dynamical models and inference from homotypic growth curves

We describe the exponential phase 1 dynamics as

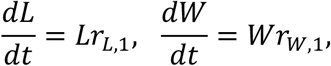

where *L* (loser) and *W* (winner) are the population sizes of 168 and 4T07, respectively, and *r*_*L*,1_ and *r*_*W*,1_ are the respective growth rates.

In phase 2, we assume a density-dependent competitive Lotka-Volterra model, parameterized in terms of intrinsic growth rates *r*_*L*,2_ and *r*_*W*,2_ and interaction terms *a*, *b*, *c* and *d*:

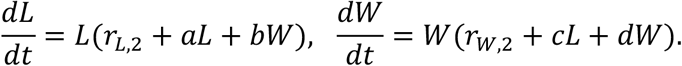

In the homotypic case, terms *bW* and *cL* vanish and the phase 2 model is equivalent to logistic growth. We combine the two models and fit to the normalized time-lapse data for the homotypic growth curves using least-squares with R package deSolve (Soetaert et al., 2010) to infer the values of *r*_*L*,1_, *r*_*W*,1_, *r*_*L*,2_, *r*_*W*,2_, *a* and *d*.

In phase 3, we assume the same model as in phase 2 except we replace *r*_*W*,2_ by *r*_*W*,3_ to account for the change in the 4T07 net growth rate (equivalent to adding a density-dependent death rate).

### Inferring between-type interaction terms

To infer the interaction parameters *b* and *c* we need data that covers a wide range of proportions of the two cell types. Since our time-lapse data is limited to only a few initial conditions, we fit the model to the outcomes of serial competition assays, and we employ the heterotypic time-lapse data for validation only. First we define

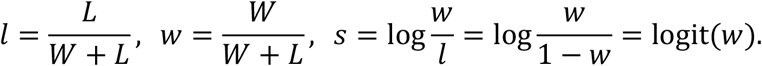

The time derivative of the *s* is then equal to the net growth rate difference, which in phase 2 is

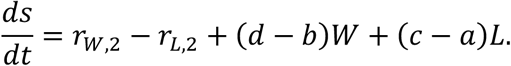

In the limit *w* → 1, the final term (*c* − *a*)*L* is negligible and we can obtain *b* in terms of 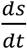, *W*, and parameters whose values we have already inferred, as follows:

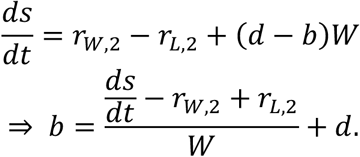

To obtain *W*, we note that in the limit *w* → 1,

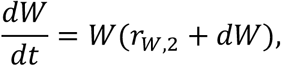

which is the logistic differential equation with solution

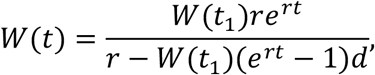

where *r* = *r*_*W*, 2_ and *t*_1_ is the time at which phase 2 begins. We can thus use our previously inferred parameter values to obtain *W*(*t*) at every time *t* in phase 2 (note that if there were not an analytical solution then we could have solved the equation numerically).

Since *W* and 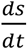 are linearly related, we can replace them by their mean values:

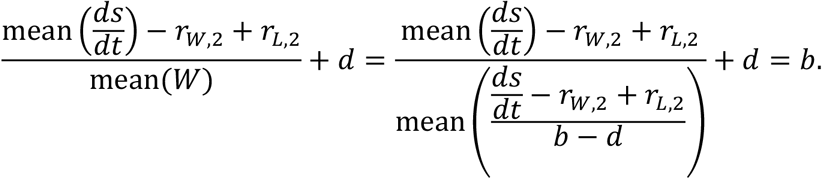

Using the mean values to calculate *b* is convenient as our competition assays reveal only the initial and final values of *s*. Specifically, we take the means in the interval [*t*_1_, *t*_2_], where *t*_2_ is the time at which phase 2 ends and

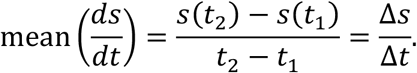

It remains only to obtain the value of the above expression – known as the gain function – in the limit *w*(*t*_1_) → 1. From competition assay data, we can immediately obtain 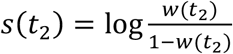 for each value of 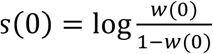. To infer *w*(*t*_1_) and *s*(*t*_1_), we need to adjust for the exponential growth of both cell types during phase 1:

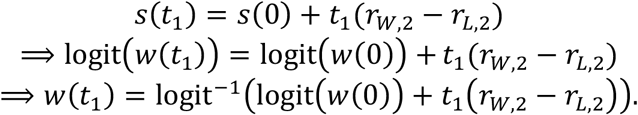

We thus obtain the values of *s*(*t*_1_) and *w*(*t*_1_) in each competition assay. Finally, we determine by linear regression the relationship between Δ*s*/Δ*t* and *w*(*t*_1_) (Supplementary figure 5B) and, from the equation of the regression line, infer the value of Δ*s*/Δ*t* in the limit *w*(*t*_1_) → 1. We then have everything required to infer the value of *b*. By an analogous method (switching *L* and *W*, *b* and *c*, and *a* and *d*) we also infer the value of *c*.

### Excluding results of first-round competition assays

In our regression to determine the relationship between Δ*s*/Δ*t* and *w*(*t*_1_), we excluded data from the first round of competition assays (days 0 to 3 in Figures 3B and 3C) because these measurements were unusually variable, and this variance was most likely an experimental artefact. Specifically, setting up the initial experiment took substantially longer than carrying out subsequent replatings as additional steps were required before seeding the cells. Since cells were kept for longer in suspension before the first round, they will have experienced more stress and potentially mortality. This means that results of the first round of competition assays are likely to be less reliable than results of subsequent rounds. For completeness, Supplementary Figures 5C and 5D show linear regression applied to the entire data set, including the first round.

### Carrying capacities

To find carrying capacities, we note that the phase 2 model can alternatively be parameterized as

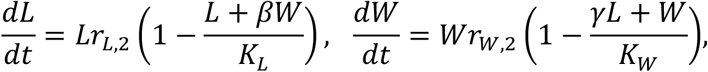

where the parameters are calculated as in Table 1. The carrying capacities *K*_*W*_ and *K*_*L*_ are the upper limits approached by the population sizes of *W* and *L*, respectively, during phase 2.

### Potential for coexistence *in vivo*

In a growing tumor, we expect cell-cell competition to be less than in our *in vitro* experiments, because, in the former, resources are continually replenished and waste materials removed by the host circulatory system. The evolutionary dynamics will then mostly depend on the difference in intrinsic growth rates and interactions mediated by diffusible factors. Furthermore, during tumor growth, the dynamics may be better described by a frequency- rather than a density-dependent model. We can then describe the evolutionary dynamics within the framework of evolutionary game theory using the payoff matrix

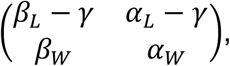

where *α*_*L*_, *α*_*W*_ < 0 denote the harm inflicted by *W* on *L* and *W*, respectively; *β*_*L*_, *β*_*W*_ > 0 are the benefits bestowed by *L* to *L* and *W*, respectively; and *γ* > 0 is the difference between the intrinsic exponential growth rates. The relative values of the entries in the payoff matrix determine which game (for example, prisoner’s dilemma or hawk-dove) is equivalent to the evolutionary dynamics.

The parameter values inferred for phase 2 of the competition assays imply

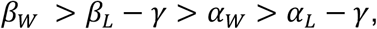

in which case the evolutionary dynamics are equivalent to a prisoner’s dilemma game for which *W* is the only evolutionarily stable strategy (ESS). This means that *W* (4T07) can invade and stably replace a population of *L* (168).

If instead *α*_*L*_ − *γ* > *α*_*W*_ then the payoff matrix defines a hawk-dove game that permits coexistence. In this scenario, *W* harms itself more than it harms *L*, and this difference outweighs *W*’s higher intrinsic growth rate. This could happen, for example, if harmful factors produced by *W* imperfectly diffuse, so that *W* cells experience a higher concentration than *L* cells. At the mixed ESS, the *W* proportion is

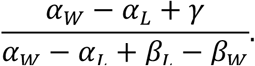

However, if additionally *β*_*L*_ − *β*_*W*_ > *γ* (so that *L* benefits itself more than it benefits *W*, and this difference outweighs *W*’s higher intrinsic growth rate) then coexistence again becomes impossible as the game again becomes a prisoner’s dilemma but with *L* as the ESS.

In a resource-poor environment, we might describe the evolutionary dynamics using the payoff matrix

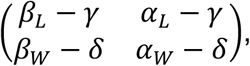

where *δ* is the reduction in *W*’s intrinsic growth rate due to the degraded environment (as inferred for phase 3 of our 96-hour competition assays). This scenario favours *L* and suggests that *L* may be the ESS in a resource-poor environment, such as hypoxic regions within a tumor.

