## Supplemental figures for "Paracrine behaviors arbitrate parasite-like interactions between tumor subclones"

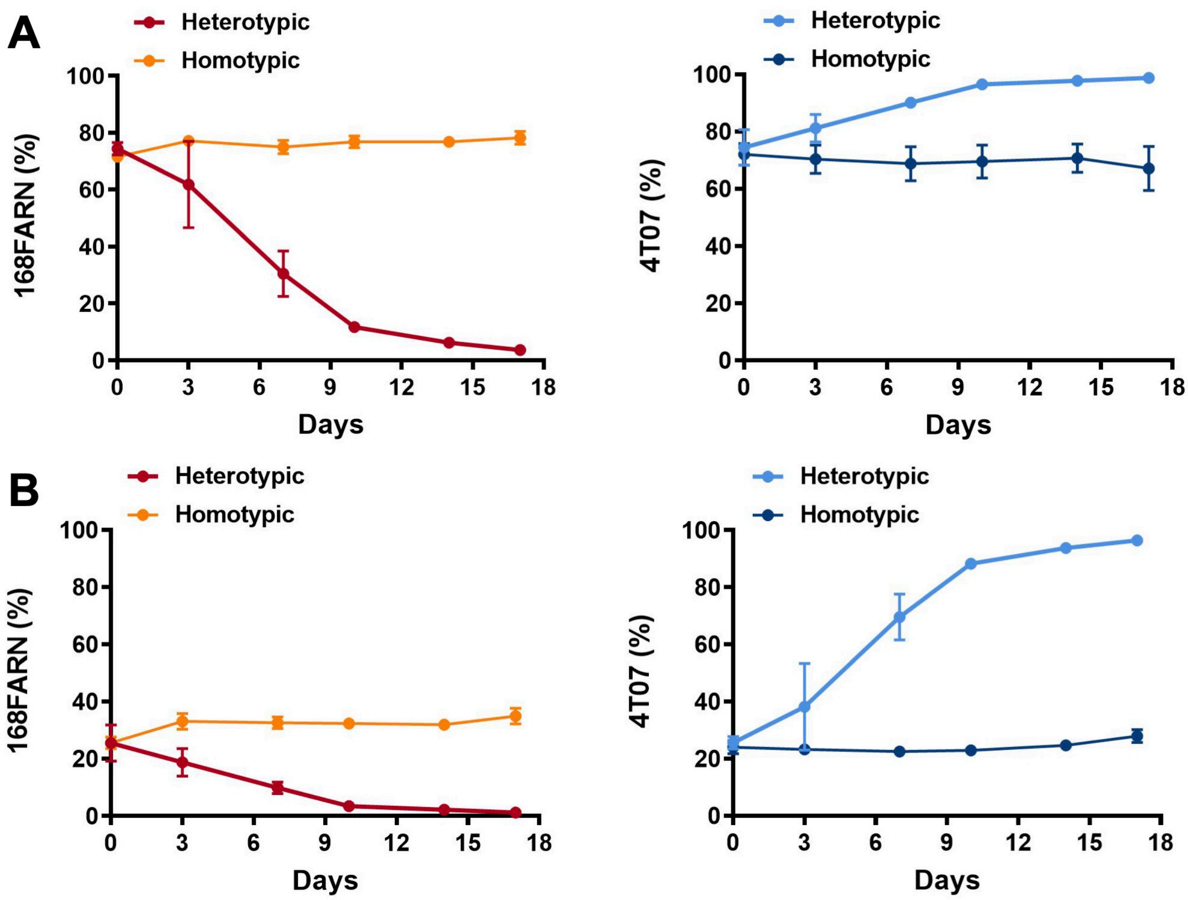


**Supplementary Figure 1**

**A and B: Growth dynamics of subclones under homotypic and heterotypic conditions.** $10^5$ cells were seeded at a 3:1 (A) or 1:4 (B) ratios in homotypic (parental and GFP expressing derivative of the same cell line) or heterotypic (different cell lines, one expressing GFP) co-cultures and harvested and replated at the initial densities ($10^5$ cells/plate) at indicated times. The ratios of GFP-labelled to unlabelled cells were estimated by flow cytometry. The results represent data from 3 independent experiments and are shown as mean +/- SEM.

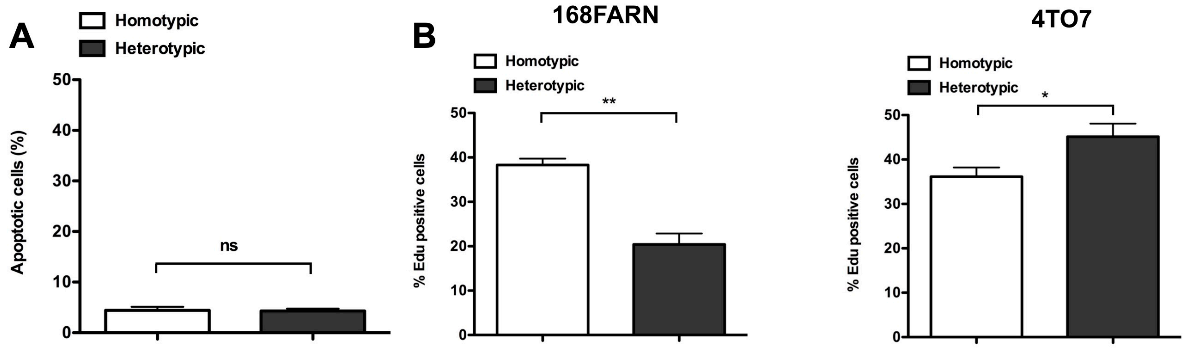


**Supplementary Figure 2**

**A: Apoptosis quantification of subclones under homotypic and heterotypic conditions.** A total of $10^5$ cells were seeded. 168G cells were co-cultured with either the 168P (homotypic) or 4T07P (heterotypic) cells at a 1:1 ratio for 4 days and harvested. Apoptosis was quantified by flow cytometry following Annexin-V staining. ns: not significant. **B: S phase quantification of subclones under homotypic and heterotypic conditions.** A total of $10^5$ cells were seeded. 168G cells were co-cultured with either the 168P (homotypic) or 4T07P (heterotypic) cells at a 1:1 ratio for 4 days. Before harvesting at day 4 cells were labelled by a 2hr pulse of EdU and the fraction of cells in the S phase was determined by flow cytometry. $^*p<0.05$, $^{**}p<0.01$

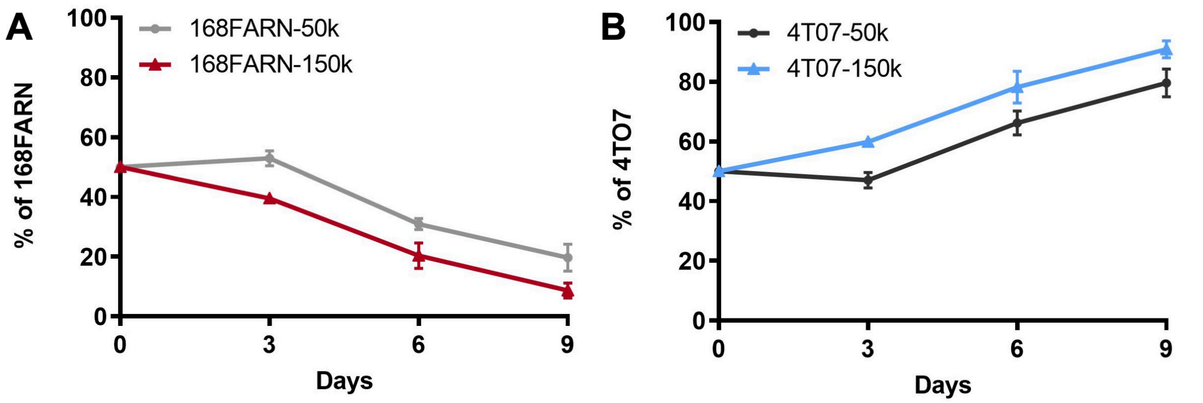


**Supplementary Figure 3**

**A: Growth dynamics of subclones at low and high density.** Experiments were performed as in Figure 2B. Cells were grown in heterotypic conditions at a starting ratio of 1:1. Cells were seeded either at low density (50K) or high density (150k), diluted and quantified every 3 days. At low density, cells do not reach confluence before replating. The results represent data from 3 independent experiments and are shown as mean +/- SEM.

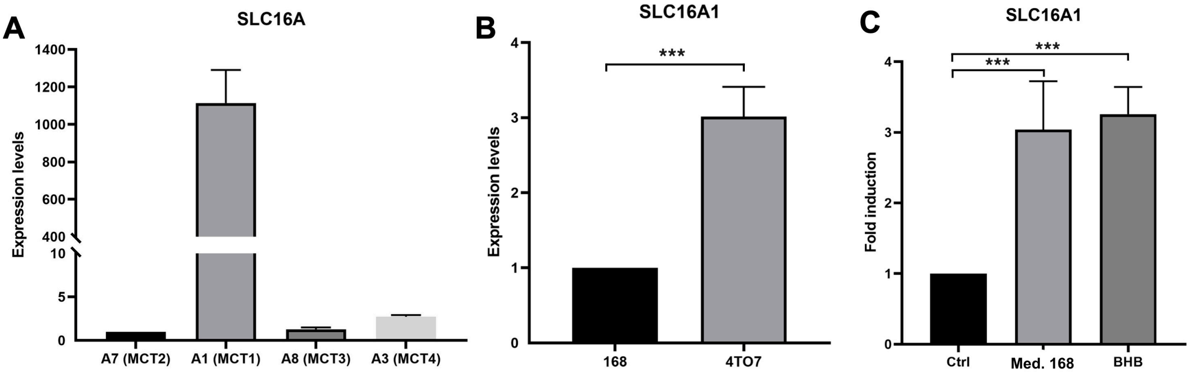


**Supplementary Figure 4**

**A:** ***Expression levels of the slc16A family transporter genes in 168FARN .*** RT-QPCR analysis was performed on 168FARN RNA for Mct2, Mct1, Mct3 and Mct4 genes and normalized to HPRT. Relative expression levels were compared to Mct2. **B: Slc16A1 expression in both subclones.** Slc16A1 RNA levels were monitored by RT-QPCR, normalized with HPRT and adjusted relative to levels in 168FARN cells cells. **C: Influence of Slc16A1 expression by β-hydroxybutyrate.** Experiment was performed as in Figure 5B. Slc16A1 RNA levels were quantified as in A and adjusted relative to levels in control condition. ***p<0.001

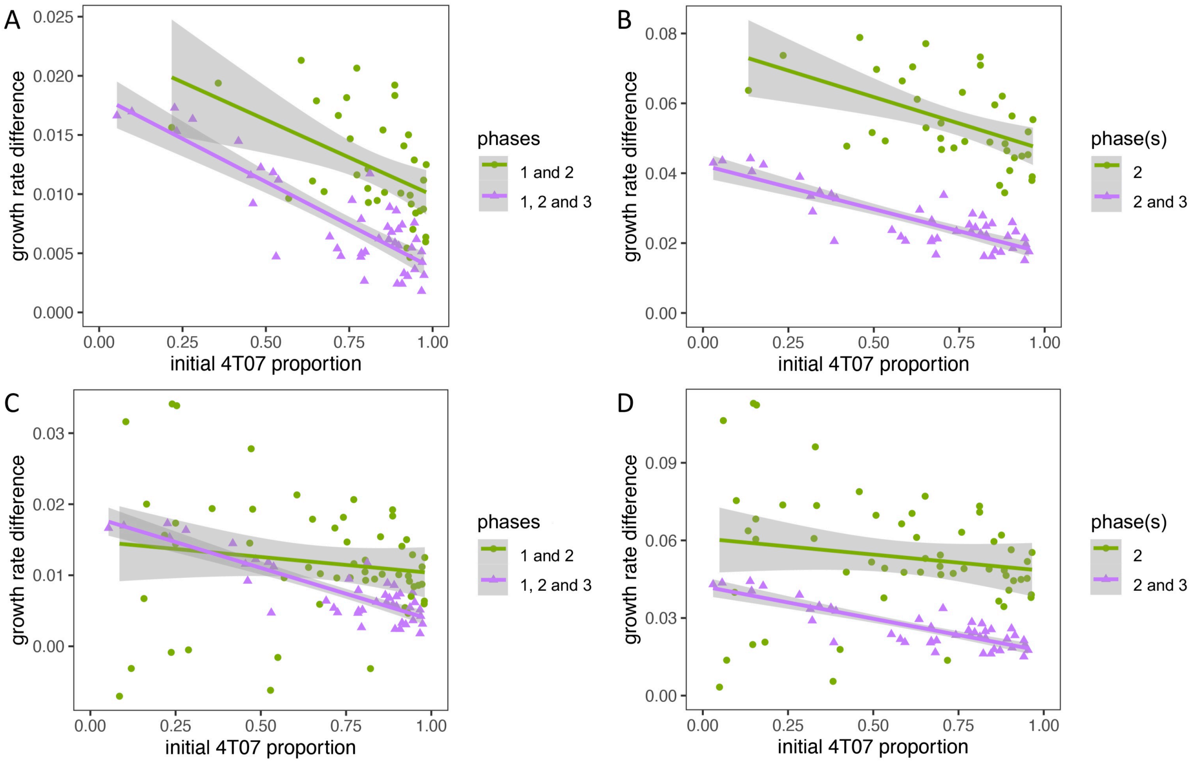


**Supplementary figure 5.**

**A: Mean net growth rate difference (gain function) versus initial 4T07 proportion in phases 1 and 2 (purple) and phases 1, 2 and 3 (green).** Each point corresponds to the outcome of a competition assay. Regression lines are shown with 95% confidence intervals. **B: Mean net growth rate difference versus initial 4T07 proportion in phase 2 (purple) and phases 2 and 3 (green)**. This data set was obtained from the data shown in **A** by adjusting for exponential growth in phase 1 (see Methods). **C: The same as A but including results for the first round of competition assays (days 0-3).** First-round measurements were excluded from analyses as they were unusually variable and unreliable due to an experimental artefact (see Methods). **D: The same as B but including results for the first round of competition assays (days 0-3).**

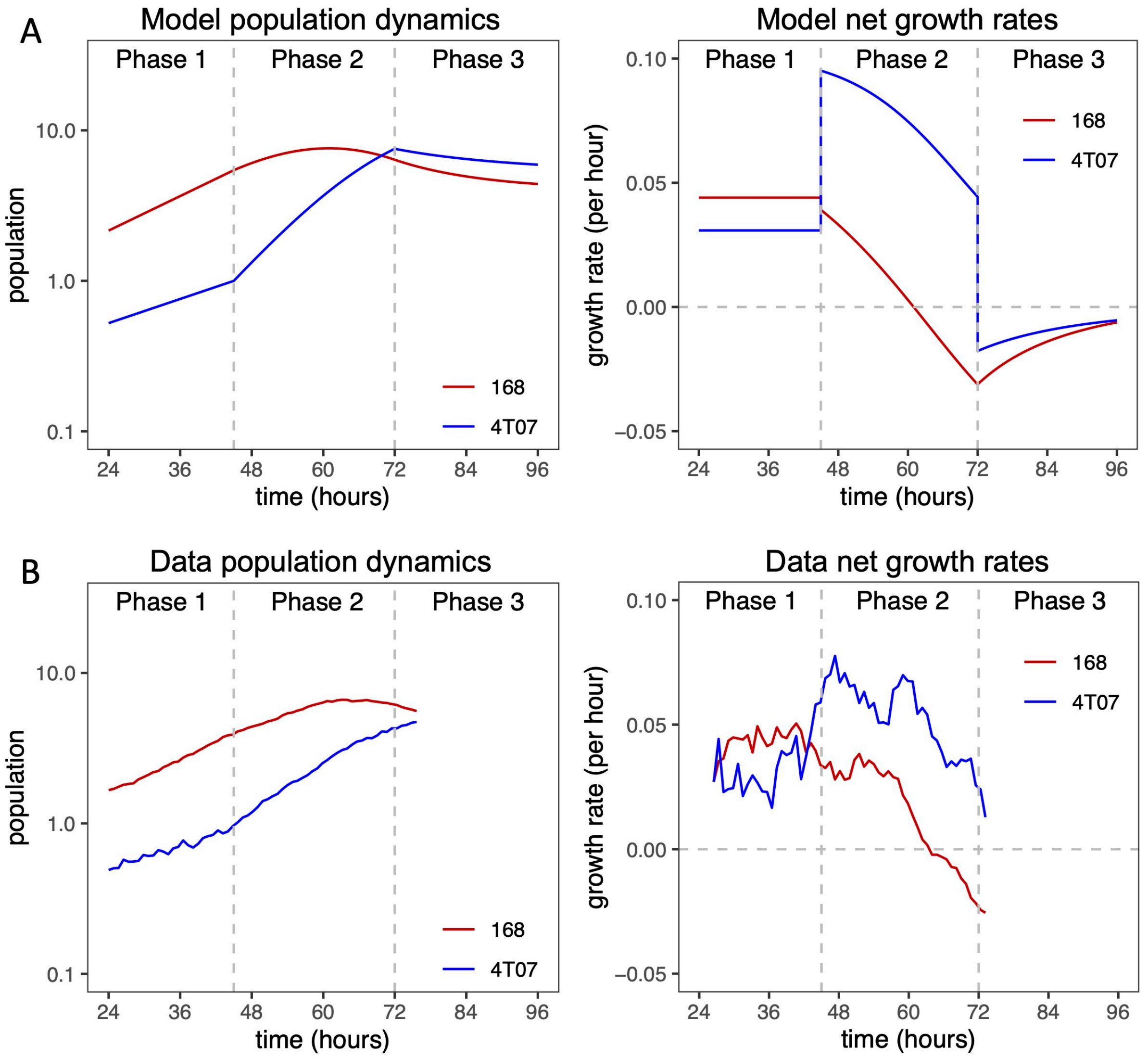


**Supplementary figure 6. Relationship between population dynamics and net growth rates.**

The net growth rate of each cell type (right column) is the derivative of its log-transformed growth curve (left column). **A: Mathematical model dynamics.** From the dynamical model, net growth rates can be found precisely by evaluating differential equation terms. The model was parameterized with values inferred from data (Table 1) and initiated with a 3:1 ratio of 168 to 4T07. **B: Empirical dynamics.** From time-lapse data, net growth rates can be approximated as local gradients (difference quotients). In this example, we estimated net growth rates from smoothed growth curves by calculating difference quotients across a 5-hour span. Smoothed growth curves (not shown) were obtained by computing running medians with a 5-hour span. Since we did not use heterotypic time-lapse data for parameter inference, the resemblance between the two rows of this figure contributes to validating our model. The data in **B** is the same as in Figure 2A and 2B.

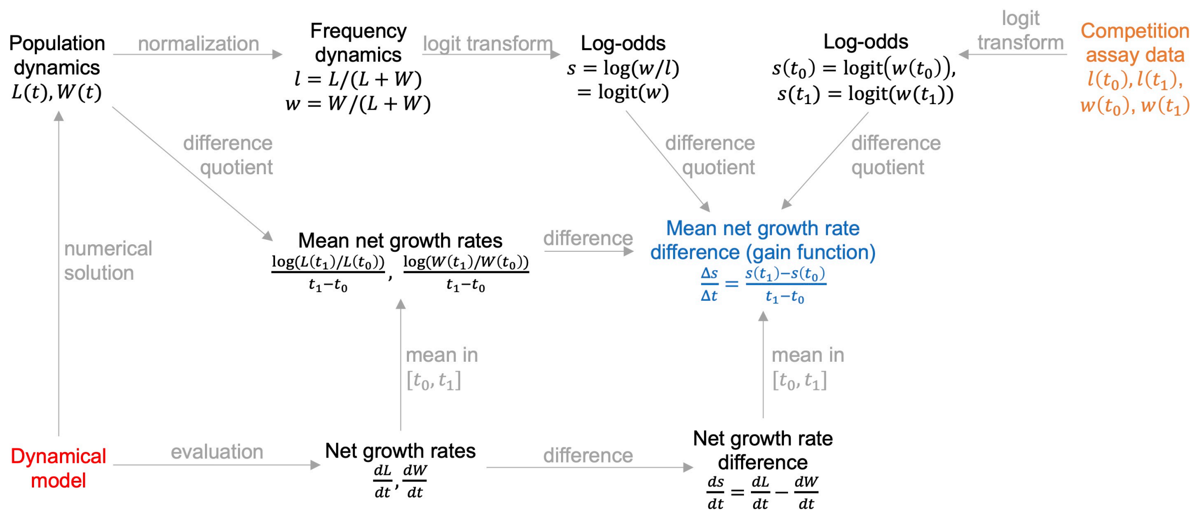


**Supplementary figure 7. Mathematical relationships relevant to our methods.**

The diagram illustrates several equivalent ways of calculating the mean growth rate difference (gain function, blue) from the parameterized dynamical model (red). Also shown is our method of calculating the gain function from competition assay data (orange).
